# CryoEM-enabled visual proteomics reveals *de novo* structures of oligomeric protein complexes from *Azotobacter vinelandii*

**DOI:** 10.1101/2025.02.04.636493

**Authors:** Rebeccah A. Warmack, Ailiena O. Maggiolo, Yuanbo Shen, Tianzheng Zhang

**Affiliations:** Division of Chemistry and Chemical Engineering 147-75 California Institute of Technology Pasadena, California 91125, United States

**Keywords:** Visual proteomics, nitrogen fixation, anerobic, cryoEM, cryoET

## Abstract

Single particle cryoelectron microscopy (cryoEM) and cryoelectron tomography (cryoET) are powerful methods for unveiling unique and functionally relevant structural states. Aided by mass spectrometry and machine learning, they promise to facilitate the visual exploration of proteomes. Leveraging visual proteomics, we interrogate structures isolated from a complex cellular milieu by cryoEM to identify and classify molecular structures and complexes *de novo*. That approach determines the identity of six distinct oligomeric protein complexes from partially purified extracts of *Azotobacter vinelandii* using both anaerobic and aerobic cryoEM. Identification of the first unknown species, phosphoglucoisomerase (Pgi1), is achieved by comparing three automated model building programs: CryoID, DeepTracer, and ModelAngelo with or without *a priori* proteomics data. All three programs identify the Pgi1 protein, revealed to be in a new decameric state, as well as additional globular structures identified as glutamine synthetase (GlnA) and bacterioferritin (Bfr). Large filamentous assemblies are observed in tomograms reconstructed from cryoFIB milled lamellae of nitrogen-fixing *A. vinelandii*. Enrichment of these species from the cells by centrifugation allows for structure determination of three distinct filament types by helical reconstruction methods: the Type 6 Secretion System non-contractile sheath tube (TssC), a novel filamentous form of the soluble pyridine transhydrogenase (SthA), and the flagellar filament (FliC). The multimeric states of Pgi1 and SthA stand out in contrast to known crystallographic structures and offer a new structural framework from which to evaluate their activities. Overall, by allowing the study of near-native oligomeric protein states, cryoEM-enabled visual proteomics reveals novel structures that correspond to relevant species observed *in situ*.

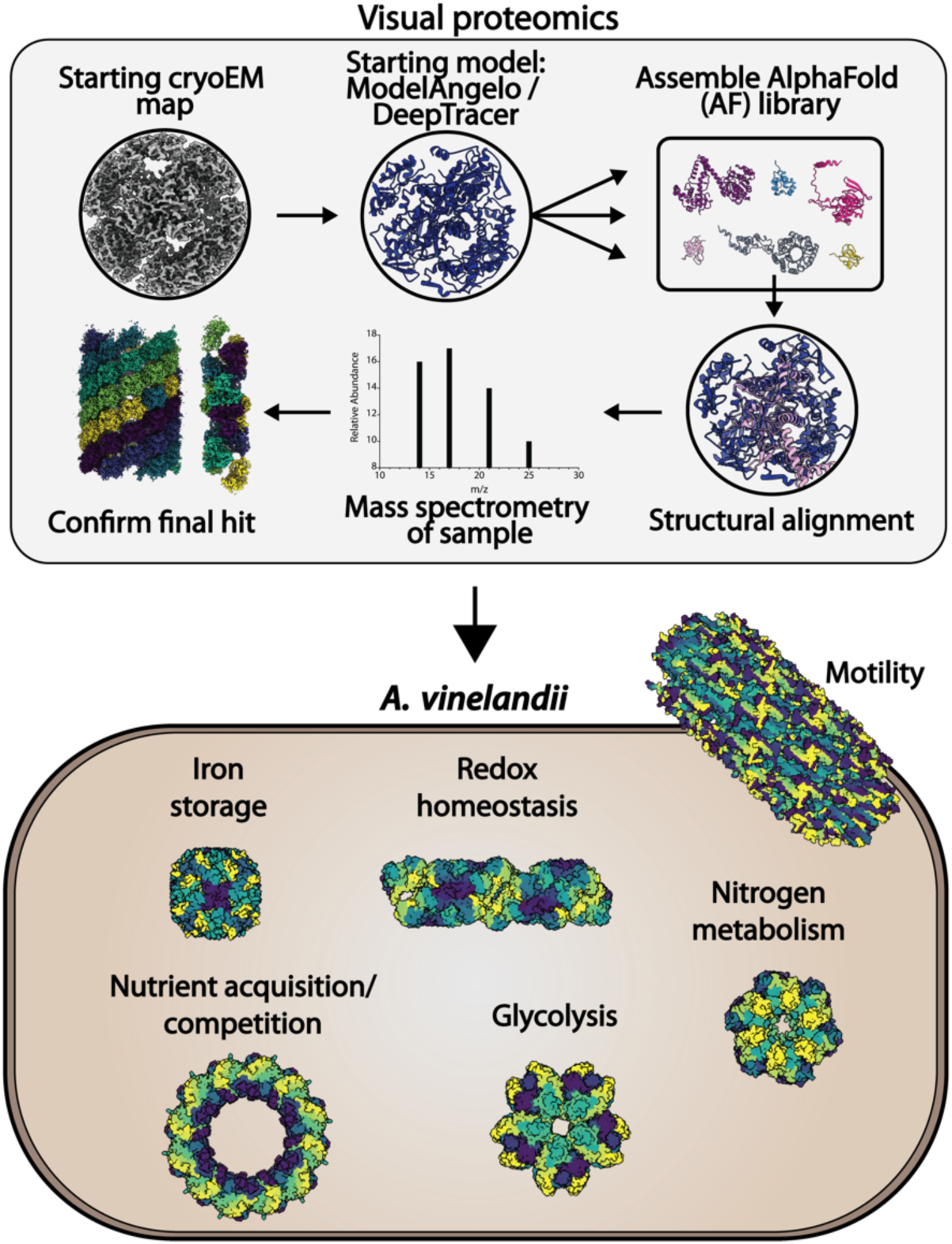

## Introduction

Emerging cryoEM methodologies continue to extend the reach of structural biology, not only facilitating the resolution of multiple conformational states, but also accommodating greater heterogeneity in sample preparations^1,2^. This opens the door for identification of unexpected protein interaction partners, interactions with cellular ultra-structures, and the identification of novel protein structures from complex mixtures. These advances are critically enabled by cryoEM processing algorithms that permit the classification of distinct species from a single dataset^3,4^, and between different conformational states of a single protein^5^. Emerging computational methods have further allowed the unambiguous identification of minor protein species or proteins in partially purified cell lysates or heterogeneous mixtures^6,7^. Collectively, these methods are steps toward the visual identification of protein molecules directly from 3D electrostatic potential maps obtained from single particle reconstructions or subtomogram averaging.

Machine learning approaches have further enabled the *de novo* determination of protein structures, with or without sequence inputs^8,9,10^. They have enabled the use of automatically built models to identify unknown proteins from cryoEM maps by comparison to experimental structures in the Protein Data Bank (PDB) or the now extensive database of AlphaFold-predicted protein structures^9,11,12^. Using similar approaches, we previously reported the application of DeepTracer coupled with AlphaFold and proteomics for the identification of an unknown 15 kDa protein as part of a nitrogenase enzyme complex^13^. The structure was derived from a single particle cryoEM dataset of the nitrogenase MoFe-protein purified from lysates of a *ΔnifV* strain of the nitrogen-fixing (diazotrophic) *Azotobacter vinelandii* bacterium. That discovery inspired the present interrogation of protein species observed in images of partially purified *A. vinelandii* extracts.

We now combine proteomic analysis of *A. vinelandii* extracts with cryoEM models built into a < 4 Å resolution maps by the automated model building programs, CryoID, DeepTracer, or ModelAngelo. The approach identifies a novel decameric state of phosphoglucoisomerase (Pgi1) and five other unknown species present in two additional datasets, identified from maps ranging in resolution from 1.9 Å – 3.7 Å: glutamine synthetase (GlnA), bacterioferritin (Bfr), and filaments of the Type 6 Secretion System non-contractile sheath tube (TssC), the soluble pyridine transhydrogenase (SthA), and the flagellum (FliC). Importantly, we also demonstrate that in the absence of prior proteomic information, species can be identified by structural alignment of DeepTracer models with AlphaFold models of *A. vinelandii* proteins. The GlnA, BfR, TssC, and FliC structures confirm the successful application of these workflows to known oligomeric structures, while the Pgi1 and SthA structures reveal striking new multimer states as a pentamer of dimers, and a filament respectively. The filamentous state of SthA further suggests that the helical ultrastructure may be used as a regulatory mechanism controlling the metabolic pools of NADH and NADPH^14^. Overall, cryoEM-enabled visual proteomics is a promising approach for the study of complex protein mixtures, and we now show it can reveal significant new insights into the physiologically relevant states of important metabolic enzymes.

## Results

### Evaluation of visual proteomic pipelines for *de novo* determination of protein structures in *A. vinelandii*

The structure of the endogenously expressed nitrogenase MoFe-protein isolated from a strain of *A. vinelandii* with the homocitrate synthase gene *nifV* disrupted^15^ was previously published by our group (MoFe^ΔnifV^)^16^. The oxygen-sensitive target protein was isolated by a two-step air-fre cd /central/groups/reeslab/rwarmack/software/cryosparc e purification comprised of anion exchange followed by size exclusion chromatography; a proteomic analysis of the purified sample was conducted as part of this study and included with the publication of the anaerobic MoFe^ΔnifV^ structure. Unexpectedly, 2D classification steps during cryoEM data processing of this same dataset revealed an abundant five-fold symmetric species (**Fig. 1A-B**) of unknown identity. Subsequent targeted processing of this species yielded a 2.7 Å resolution reconstruction without imposed symmetry (**Fig. 1C, left panel; Supp. Fig. 1**). Visual analysis of the cryoEM map suggested a decameric protein arranged as a pentamer of dimers. Correspondingly, enforcement of D5 symmetry during refinement yielded a 2.5 Å resolution reconstruction (**Fig. 1C, right panel**).

**Figure 1:**
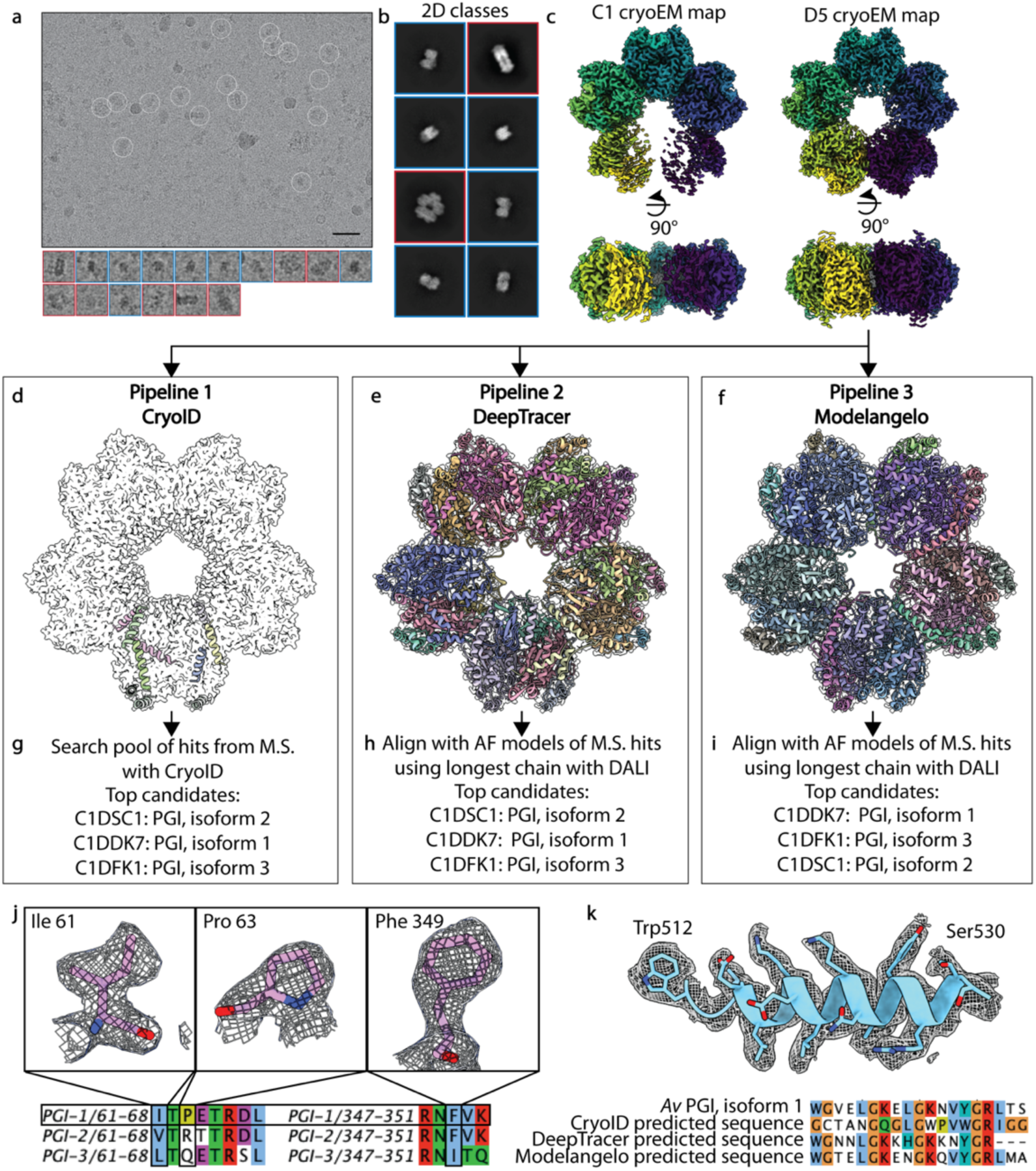
Identification of an unknown *A. vinelandii* protein map from single particle cryoEM as phosphoglucoisomerase (Pgi). A, Representative micrograph from single particle dataset. Scale bar represents 50 nm. Circles and insets correspond to extracted particles. Particles boxed in blue represent the expected nitrogenase MoFe-protein. Particles boxed in red represent the unknown protein. B, Representative 2D classes from cryoEM data processing. 2D classes boxed in blue represent the expected nitrogenase MoFe-protein. 2D classes boxed in red represent the unknown map. C, Left panel, 2.7 Å resolution reconstruction of abundant species with no symmetry imposed. Right panel, 2.5 Å resolution reconstruction of abundant species enforcing D5 symmetry. D-F, computational pipelines for the identification of the unknown protein using the available softwares CryoID, DeepTracer, and ModelAngelo, respectively. G-I, Top three hits corresponding to the unknown species using the respective pipelines. J, Identification of the unknown contaminant as Pgi1 over isoform 2 and 3. Upper panel, density corresponding to residues Ile 61, Pro 63, and Phe 349 in the refined structure. Lower panel, sequence alignment of *A. vinelandii* Pgi isoforms 1, 2, and 3 color-coded according to Clustal. K, Upper panel, segmentsfit into cryoEM density built by ModelAngelo. Lower panel, sequence alignment of the illustrated segment and *A. vinelandii* Pgi, isoform 1.

Three general workflows were evaluated as discovery tools for the identification of this species (**Fig. 1D-I**): First, the CryoID program was employed against the D5-symmetry enforced cryoEM map, with the proteomic results as inputs (**Fig. 1D**)^8^. Second, the DeepTracer program was run inputting only the D5-symmetry enforced cryoEM map as an input without any sequence information (**Fig. 1E**)^9^. Third, the ModelAngelo program was run using only the D5-symmetry enforced cryoEM map as an input and no sequence information (**Fig. 1F**)^10^. In parallel, a library was generated of the AlphaFold structures of 520 proteins identified by bottom-up proteomics analysis. Subsequently, the resulting models from DeepTracer and ModelAngelo were aligned to each of these AlphaFold predicted models using the program DALI^17^, while the CryoID segments were sequentially compared to the proteomic results with the search pool function of the program^18,19^.

All three pipelines identified the same top three hits as isoforms of the glycolytic enzyme phosphoglucosoisomerase (accession codes C1DSC1, C1DDK7, and C1DFK1), although in varying orders (**Fig. 1G-I**). The C1DDK7 (Pgi1), C1DSC1 (Pgi2), and C1DFK1 (Pgi3) species were identified at 6.5%, 0.01%, and 0.24% relative abundance in the original proteomic analysis, respectively^16^ (**Supplementary Table 2**). As pipelines 1-3 identified all three *A. vinelandii* Pgi isoforms as top hits, they were compared to cryoEM densities of the higher resolution D5-symmetry enforced map, with particular emphasis on regions of high sequence divergence, to determine if a specific isoform could be verified (**Fig. 1J**). Several regions definitively matched the Pgi1 isoform, namely the densities for Ile61, Pro63, and Phe349, suggesting that this structure is representative of the Pgi1 isoform. Additionally, sequence prediction by ModelAngelo based on the cryoEM density matched the sequence of Pgi isoform 1 with ∼77% sequence identity (**Fig. 1K**). However, as all three isoforms were identified by the proteomic analysis, we could not rule out that this structure represents an average across all three forms. Independent of the proteomic results, DeepTracer and ModelAngelo models were also compared against the Protein Data Bank (PDB) using the DALI server^17^, and the top hits were PDB 4QFH (*T. cruzi* Pgi)^11^ and PDB 3NBU (*E.coli* Pgi)^20^, respectively; further confirming the identified structure as Pgi. Side chains were added to the DeepTracer model corresponding to the Pgi1 sequence using SCRWL4^21^ and ModelAngelo was re-run using this sequence as an input. The subsequent models were assessed for map-to-model fit using Q-scores. The DeepTracer model demonstrated an average Q-score of 0.42, while the ModelAngelo model showed an average Q-score of 0.70 (**Supp. Fig. 2**)^22^. Thus, the ModelAngelo results were used as a starting model for subsequent structure building and refinement.

### The *A. vinelandii* Pgi1 structure exhibits distinct conformational states

The decameric Pgi1 structure contains two dominant interfaces, each created by a pair of Pgi1 monomers. Interface A buries ∼5950 Å^2^ surface area between monomers in dimers related by two-fold symmetry axes. Interface B buries ∼1080 Å^2^ surface area between dimers related by the five-fold rotation axis (**Fig. 2A**). A comparison of Pgi1 sequences across several model organisms indicated that residues involved in Interface A are 45% identical, while residues involved in Interface B are 18% identical, with the strongest sequence similarity occurring between *A. vinelandii* and *Pseudomonas aeruginosa* (**Supp. Fig. 3**). AlphaFold predictions containing five copies of the Pgi1 chain were able to replicate Interface B in the *A. vinelandii* and *P. aeruginosa* models, but did not form Interface B in any of the other structures, perhaps suggesting that this oligomeric state may be limited to bacteria.

**Figure 2:**
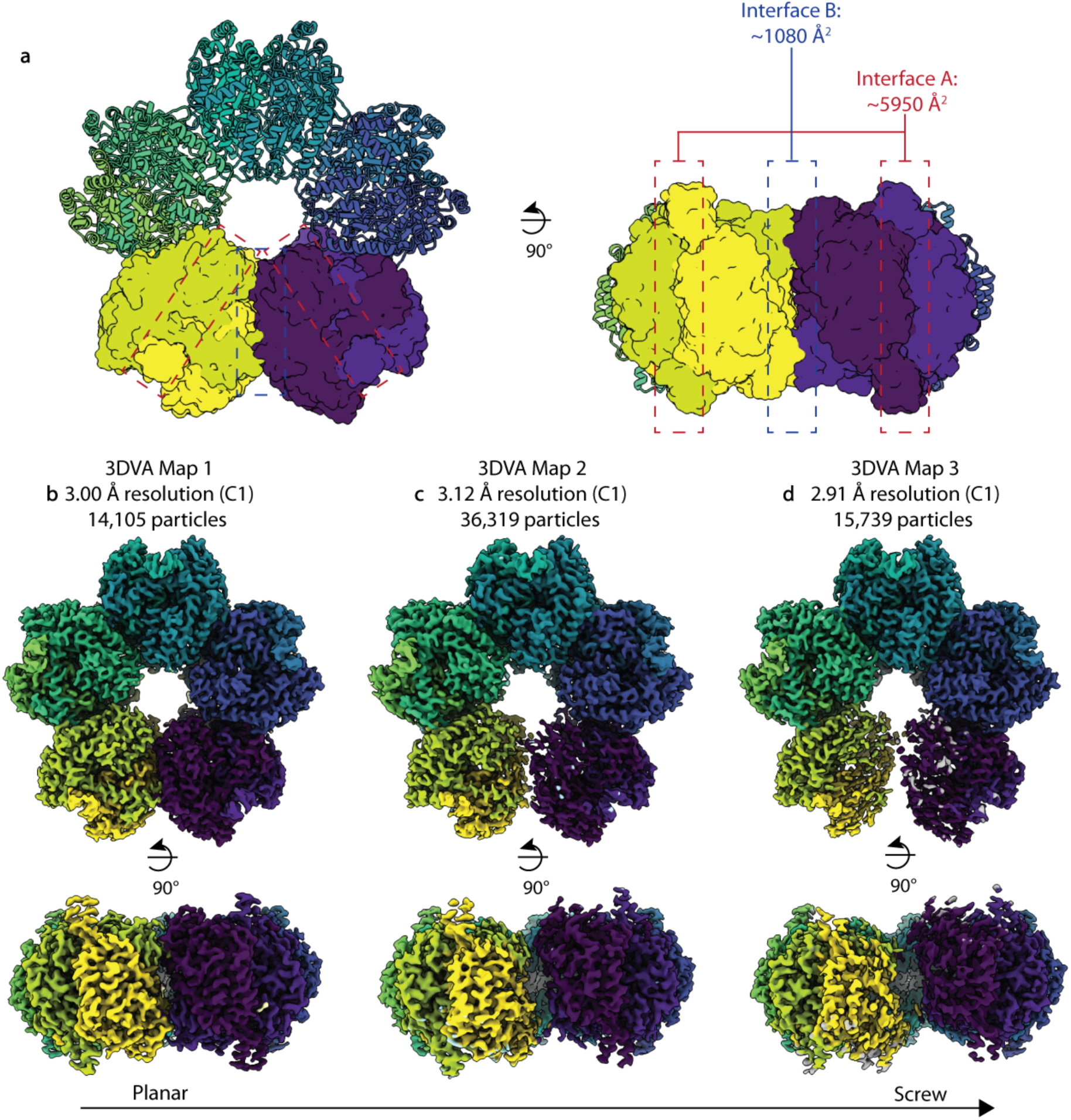
The *A. vinelandii* Pgi1 exhibits distinct conformational states. A, Refined structure of Pgi1 into D5 single particle cryoEM map. The upper three dimers are shown as cartoon representations, the lower two dimers are shown as surface representations. The intradimeric interfaces are indicated with a dashed red box, the interdimeric interfaces are indicated with a dashed blue box. B-D, Single particle cryoEM maps reconstructed after subclassification by three dimensional variability analysis (3DVA). These maps trend from a more planar structure along the horizontal axis to a screw structure with translation of dimers upwards and downwards.

Weaker density was observed for two of the dimers when refining the particles without imposing any symmetry constraints (C1 refined). Three dimensional variability analysis (3DVA) was performed on the subset of particles used for the C1 refinement (**Fig. 2B-D; Supp. Fig. 1**). The resulting 3DVA maps showed a progressive conformational shift, out of plane with respect to the five-fold symmetry axis, to generate a right-handed screw (**Supplementary Video 1**). Three separate maps were isolated from this particle subset using 3DVA and were reconstructed with no symmetry imposed (C1; **Fig. 2B-D**). The largest translational shift appears to occur along one Interface B in the pentameric structure (**Fig. 2D**). While contacts between the dimers remained largely constant, two monomers within the ‘screw’ state have lost significant density suggesting increased flexibility and/or partial occupancy (**Fig. 2D**). The loss of density in these two monomers, which previously formed contacts with adjacent dimers may create changes across the oligomer.

### *A priori* proteomic information is not necessary for accurate protein identification

While examining the micrographs of a sample prepared from *A. vinelandii*^23^ expressing a MoFe-protein variant, we observed an enriched, unidentified 6-fold symmetric species, from which we obtained a 3.18 Å resolution map (**Fig. 3A-C**; Supp. Fig. 1). We assembled a directory containing all *A. vinelandii* structure predictions currently within the AlphaFold3 database (totaling 4,968 proteins), then generated a DeepTracer model from the cryoEM map, and finally compared this model to the *A. vinelandii* AlphaFold structure predictions using DALI structural alignment (**Fig. 3D**). The top alignment for the longest chain in the DeepTracer model was glutamine synthetase, which synthesizes glutamine from glutamate and ammonia (**Fig. 3E**; GlnA; **Supplemental Table 3**). Subsequent proteomic analysis of the protein sample confirmed the GlnA protein is present in the sample (**Supplemental Table 3**). The DeepTracer model was then compared to the PDB using the DALI server, and the top match was glutamine synthetase (**Supplemental Table 3**). This first structure of *A. vinelandii* GlnA closely resembles those of its dodecameric bacterial homologues^24^, forming 12 active sites per complex, each at the interface between adjacent monomers. The ability to identify targets *de novo* without relying on a predefined list of mass spectrometry-identified targets, demonstrates the broader applicability of this workflow.

**Figure 3:**
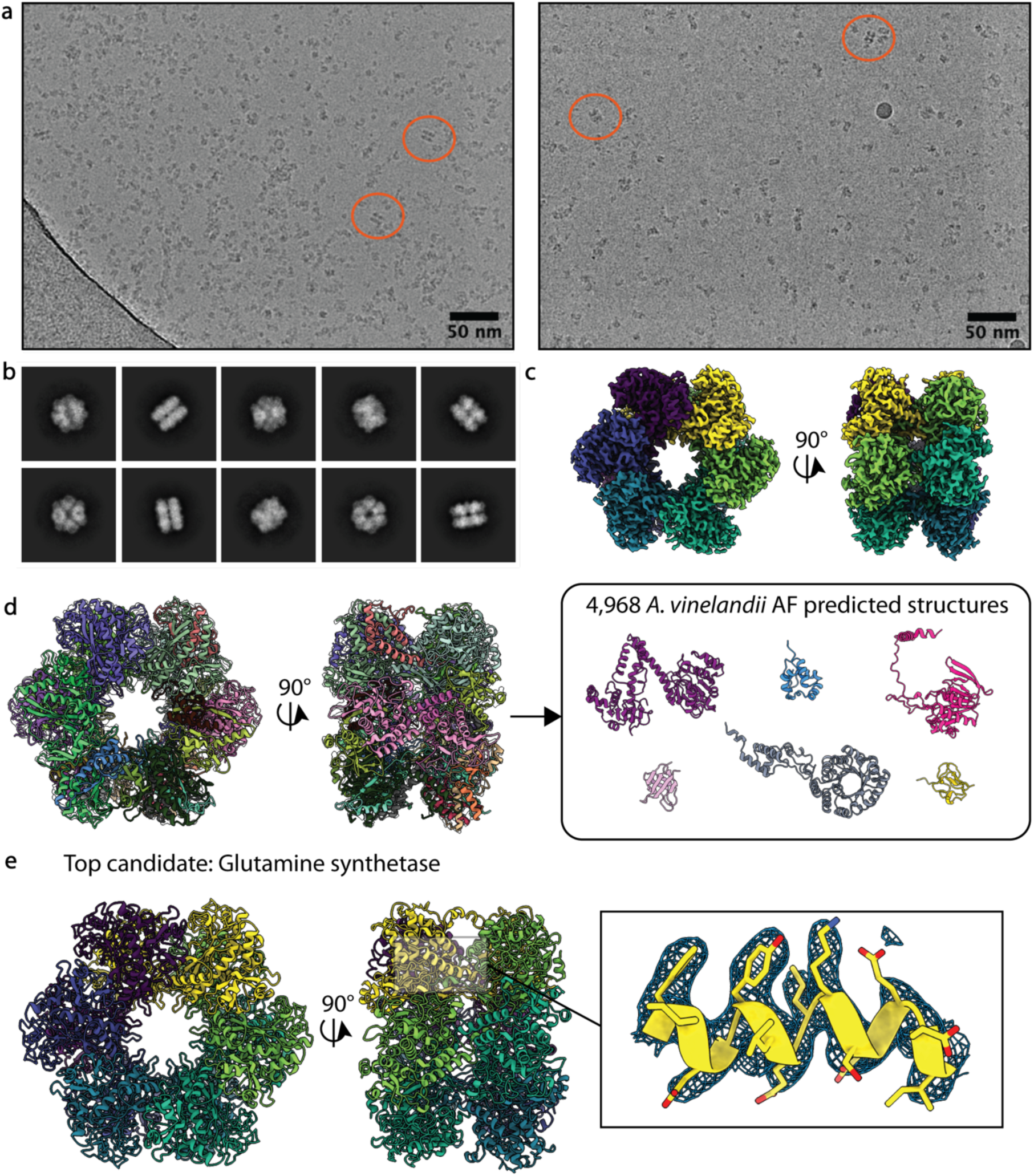
Identification of glutamine synthetase without *a priori* proteomic information. A, Representative micrographs of a MoFe^Anc^^1^ dataset containing an unknown protein (orange circles). B, 2D classes of unknown protein from panel A. C, 3.18 Å resolution reconstruction of the unknown protein in panels A and B. D, A DeepTracer model was built into the cryoEM density and structurally aligned to a directory containing all 4,968 AlphaFold models of *A. vinelandii* proteins. E, Through the pipeline in panel D, the unknown protein was identified as glutamine synthetase (GlnA).

### Application of visual proteomics to filamentous species

CryoET of intact cells allows for the study of protein assemblies in their native environment, but identification of the features observed within cells can prove challenging. To demonstrate the applicability of visual proteomics pipelines to *in situ* studies, cryoET data was collected on nitrogen-fixing *A. vinelandii* cells, revealing large filamentous assemblies within the cytoplasm (**Fig. 4A-B**). We aerobically pelleted large proteinaceous species from *A. vinelandii* grown under nitrogen-limiting conditions and analyzed the resulting sample by single particle cryoEM. Micrographs revealed a mixture of three apparently distinct filamentous species, labeled Filaments A, B, and C; as well as globular particles subsequently identified as bacterioferritin using the DeepTracer pipeline (**Fig. 4C-G; Supp. Fig. 4**). Filamentous assemblies require an accurate estimate of the helical parameters during reconstruction which can sometimes be challenging to determine. Using the helical reconstruction pipeline in cryoSPARC, and the symmetry search utility to estimate helical parameters, we were able to achieve reconstructions of Filaments A and B at nominal resolutions of 1.85 Å and 3.70 Å, respectively (**Fig. 4D-G; Supp. Fig. 1**). Comparison of micrographs containing Filament C to previous literature, coupled with proteomic results, suggested the structure belonged to flagellin (FliC)^25^, with the flagellar hook visible in many micrographs (**Supp. Fig. 4; Supplemental Table 4**). Enforcement of canonical helical parameters for flagellin resulted in ordered inner domains, but disordered outer domains (**Supp. Fig. 4**). Doubling of these helical parameters improved the resolution of the outer domains and resulted in maps with a nominal resolution of 2.82 Å, presenting a structure highly similar to the *E. coli* flagellar structure^26^ (PDB 7SN4; **Supp. Fig. 4**).

**Figure 4:**
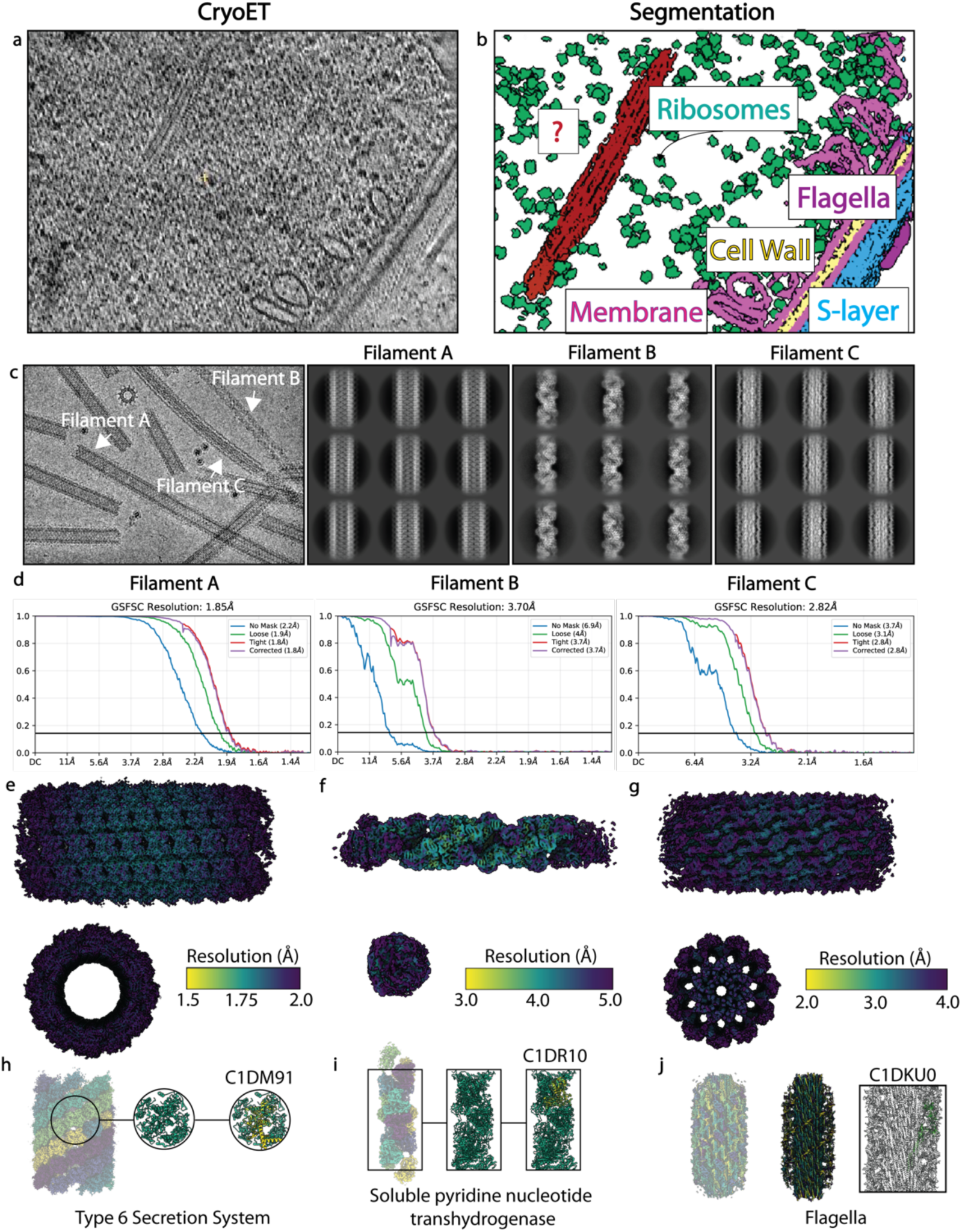
Identification of unknown filamentous species using pipeline 2. A, Tomogram of *A. vinelandii* cells under nitrogen-fixing conditions. B, Corresponding segmentation of the tomogram shown in panel A. C, Left, representative micrograph containing Filament A from a single particle dataset. Right, 2D classes of Filament A-C. D, FSC curves for Filaments A-C estimating nominal resolution. E-G, CryoEM densities for Filaments A-C, respectively, color coded according to local resolution using the viridis color palette. H-J, Identification of the following filaments using visual proteomics: Filament A (Type 6 Secretion System); Filament B (Soluble Pyridine Nucleotide Transhydrogenase); Filament C (Flagella).

Segments of the unknown Filament A and B maps were extracted using ChimeraX and DeepTracer models were built into these subvolumes (**Fig. 4H-J**). Comparison of the Filament A model to our compiled *A. vinelandii* AlphaFold database revealed the top match as the DUF877 family protein, also known as the Type 6 Secretion System non-contractile sheath tube protein TssC, which was confirmed to be present in the sample via mass spectrometry (**Supplemental Table 4**). This secretion system is generally associated with interbacterial competition through the delivery of toxic effector proteins to adjacent cells^27^.

Despite the lower resolution of the Filament B map (3.70 Å), DeepTracer was able to build a model, and comparison to our *A. vinelandii* AlphaFold database revealed the top match as the soluble pyridine nucleotide transhydrogenase (SthA), also known as an NAD(P)+ transhydrogenase (*Re/Si*-specific) (SthA; **Supplemental Table 4**). This flavoprotein balances the intracellular pools of NADH and NADPH through hydride transfer^28^. SthA was identified within the mass spectrometry results, and its predicted structure matched the DeepTracer model and density for this filament (**Fig. 4I**).

Soluble transhydrogenase (SthA) from *A. vinelandii* assembles into filaments with a rise of 38 Å and a twist of -108° (**Fig. 5A**). Despite the lower resolution of the SthA cryoEM reconstruction, it shows clear density for its FAD cofactor (**Fig. 5B**). AlphaFold predictions of the SthA monomer with FAD and NAD position the NAD substrate in contact with the FAD. This configuration is consistent with other flavoproteins, such as NADH peroxidase (PDB 2NPX; **Supp. Fig. 5**)^29^; however, no density for NADH substrates was found in our cryoEM structure. Strikingly, we have found that AlphaFold is able to predict nearly an exact match for the SthA filament when provided with more copies of the monomer (**Fig. 5C**), demonstrating its impact in predicting new oligomeric states of enzymes and its potentially broad utility for visual proteomics.

**Figure 5:**
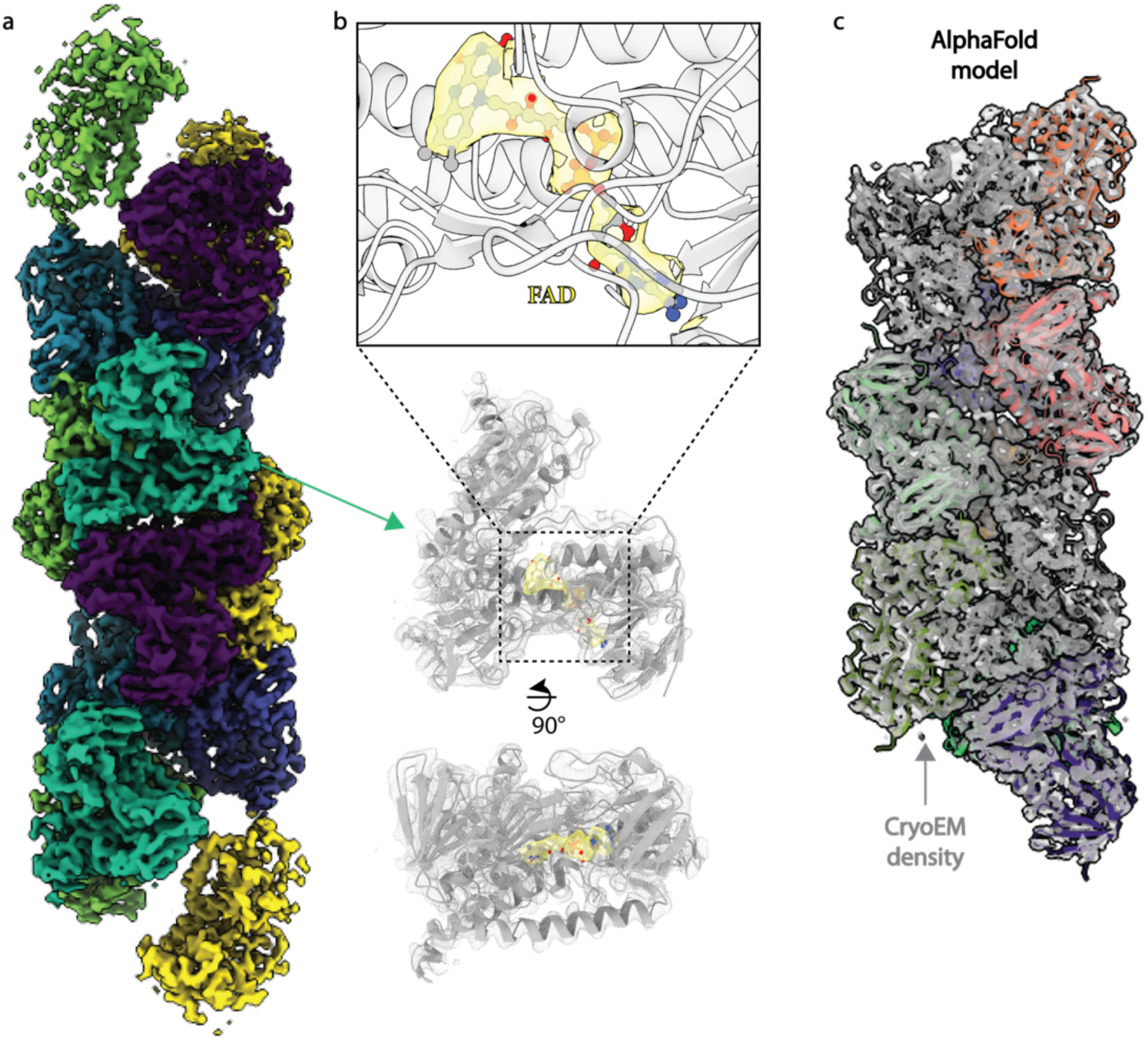
Filamentous structure of the soluble pyridine transhydrogenase. A, Overall cryoEM density for the SthA filament. B, CryoEM density for the FAD cofactor in SthA. C, AlphaFold model of 10 copies of the SthA subunit fit into the cryoEM density of the SthA filament.

## Discussion

The power of cryoEM lies in its ability to inform on the unrestrained, pseudo-solution environment of the sample, avoiding conformational selection by crystallization and harnessing algorithms that allow for robust subclassification of mixed states. Technological advancements have allowed single particle cryoEM to regularly achieve resolutions of 4 Å or better. In this study, we have compared three computational pipelines using existing software for the identification of unknown single particle cryoEM species from endogenous samples isolated from the nitrogen-fixing bacterium *A. vinelandii*. Pipeline 1 utilized the CryoID program; pipeline 2, the DeepTracer and DALI programs; and pipeline 3, the ModelAngelo and DALI programs^8–10,19^. In our primary test case all three programs identified the protein as phosphoglucoisomerase (Pgi1), which corroborated existing proteomic analyses of the sample^16^. While the program CryoID requires proteomic analysis as an input, we demonstrated that models built by ModelAngelo and DeepTracer can provide accurate identification of unknown species by structural alignment with a curated database of AlphaFold predictions. Q-score metrics applied to the ModelAngelo output models, generated with sequence information, indicated that pipeline 3 is most effective for subsequent structure refinement (**Supp. Fig. 2**). However, both DeepTracer and ModelAngelo are able to provide equivalently accurate identification of unknown structures, with web interfaces that make both programs highly accessible to new users.

Further analysis of the resulting structures from our studies reveals several surprising features. We have found the *A. vinelandii* Pgi1 assembles into a unique decameric oligomer, which has not been observed in previous crystal structures of Pgi from other organisms. The vast majority of deposited Pgi crystal structures in the PDB are dimeric, with one hexameric structure (PDB code 2Q8N)^11^. Studies in *Saccharomyces cerevisiae* have shown that six out of 10 of the glycolytic enzymes form filamentous structures, though Pgi1 was not identified as one of these enzymes^30^. It has been suggested that the oligomerization of these enzymes may provide a mechanism for regulating the flux of glycolytic intermediates into branching pathways. Indeed, the Pgi1 substrate, glucose-6-phosphate (G6P) enters the pentose phosphate pathway to generate NADPH and pentose. The discovery of higher order oligomeric states suggests that this complex of Pgi1 may play a role in controlling the partitioning of G6P between glycolysis and the pentose phosphate pathway. In *A. vinelandii* that is grown on sucrose, glycolysis is likely upregulated during nitrogen fixation, supported by the identification of three additional glycolytic proteins in our proteomic analysis^13^.

In addition to our discovery of the higher order Pgi1 oligomer, we were surprised to unearth filaments of the soluble pyridine nucleotide transhydrogenase (SthA). Membrane-bound transhydrogenases, which couple proton transfer across the membrane to NADPH cofactor transhydrogenation to form NADH, are well characterized both biochemically and structurally, however, the energy-independent soluble transhydrogenases are less well understood^31,32^. SthA is part of a larger group of flavin-dependent pyridine nucleotide dehydrogenases, and utilizes the flavin adenine dinucleotide (FAD)^32^. While membrane-associated transhydrogenases are found in bacteria and in the mitochondria of eukaryotes, SthA is only found in certain bacteria^31^. While both the *P. aeruginosa* and *A. vinelandii* SthA have been shown to oligomerize^33–35^ and form helical structures up to 1800 nm in length by negative stain electron microscopy^34,35^, no molecular structure of the assemblies has been previously reported. Importantly, although a majority of metabolic enzymes that form fibrils are active in the polymerized state^14,36^, and Broek *et al*., noted that addition of the product NADP+ could dissociated the *P. aeruginosa* oligomers observed by negative stain^34^, it is unclear whether filamentous SthA is active.

Two recent reports demonstrated the *in vitro* formation of filamentous assemblies of the *A. vinelandii* nitrogenase MoFe- and Fe-proteins with the Shetna protein II upon oxidation of the latter protein^37,38^. This protein has been reported to conformationally protect the oxygen-sensitive nitrogenase proteins under high oxygen conditions^39^. As these filaments have been associated with oxygen damage, we considered the possibility that we might also observe these filaments in our aerobically prepared pellets of Filaments A-C (**Fig. 4**). However, no such filaments were observed within either our single particle cryoEM micrographs or tomograms of the *A. vinelandii* cells. It is possible that these filaments form under higher oxygen concentrations than the ambient conditions tested here, or that other absent triggers are involved in their formation.

Each of the complexes identified here were derived from actively nitrogen-fixing *A. vinelandii* cells. While their presence in the enriched extracts does not necessarily indicate a direct link to nitrogen fixation, their integral roles in *A. vinelandii* metabolism are well understood. With this knowledge we can begin to build a basic structural network of nitrogen fixation: We begin with the reduction of dinitrogen, which requires significant levels of ATP. Pgi1 acting within the glycolytic pathway to metabolize the glucose molecules after breakdown of sucrose, generates both ATP and NADH. This NADH can enter the electron transport chain (ETC) to generate further ATP. SthA additionally ensures adequate levels of NADH for the ETC by balancing the pools of NADPH and NADH. In addition to ATP, nitrogen-fixing cells require high levels of iron for the synthesis of the nitrogenase metalloclusters which is stored by bacterioferritin. GlnA plays a central role in nitrogen metabolism by assimilating the ammonia produced by nitrogen fixation into the amino acid glutamine. In nitrogen excess conditions, the binding of glutamine to GlnA is thought to regulate repression of the nitrogenase genes^40^. The primary role of the type 6 secretion system is to export antibacterial proteins to competing microbes^27^, possibly providing *A. vinelandii* with an advantage for nutrients in challenging environments. This competitive edge for nutrients is bolstered by high motility enabled by flagella.

The successful application of the visual proteomics pipelines for the identification of unknown protein species in *A. vinelandii* extracts, and the subsequent comparison of output models offers new opportunities for *ex situ* structural analysis. As studies move towards more complex cellular contexts, the application of visual proteomics to the discovery of novel protein complexes *in situ* and *ex situ* will clarify our understanding of metabolic systems.

## Methods

### Strains

The strains used in this study are summarized in Supplementary Table 1. Strains DJ33 (ΔNifDK), DJ54 (ΔNifH), and DJ605 (ΔNifV) were a kind gift from Dr. Dennis Dean^15^. Mutated strains were generated through homologous recombination according to the protocols described by Dos Santos^41^. Briefly, plasmids were generated containing the modified proteins of interest surrounded by flanking regions with ∼500 bps homologous with the genomic site of the protein. Plasmids were transformed into *A. vinelandii* and homologous recombination was allowed to occur for 20 min at 30°C before plating onto selective media.

### Preparation of partially purified extracts with Pgi1 and GlnA

The MoFe-protein samples containing the enriched Pgi1 and GlnA species were purified as described previously^13,42^. Briefly, purifications were performed under anaerobic conditions using a combination of Schlenk line techniques and anaerobic chambers with oxygen-scrubbed argon.

### Preparation of filamentous samples

Pelleting of filamentous species was achieved using similar protocols to those described by Kudryashev *et al*^43^. Fifty mL cultures of *A. vinelandii* were harvested at ∼1.3 OD_600_ by centrifugation at 7,000 RPM for 15 min at 4°C. Cells were resuspended in 4x volumes of 25 mM Tris-HCl, pH 7.8 and 4 M glycerol. Glycerol-swollen cells were collected through an additional spin at 7,000 RPM for 15 min at 4°C. Cells were resuspended 50 mL lysis buffer (25 mM Tris-HCl, pH 7.8, 150 mM NaCl, 200 ug/mL lysozyme, 50 ug/mL DNase I, 5 mM EDTA, 0.1% SDS, and 0.5% Triton X-100). After 15 min at 37°C, 10 mM MgCl_2_ was added, and the solution was incubated for an additional 5 min at 37°C. EDTA was then added at a final concentration of 15 mM, and the solution was cleared at 10,000g for 20 min. The clarified lysate was then spun at 38,000 RPM for 1 h at 4°C in a Beckman ultracentrifuge. The resulting pellet was resuspended in 50 mL of 25 mM Tris-HCl, pH 7.8, 150 mM NaCl, 0.1% SDS, and 0.5% Triton X-100 and spun again at 38,000 RPM for 1 h at 4°C. The resulting pellet was resuspended in 50 mL of TBS and spun again at 38,000 RPM for 1 h at 4°C. The final pellet was resuspended in 500 uL of TBS, then concentrated to 50 uL in a 100 kDa MWCO spin filter. This sample was used directly for cryoEM grid preparation as described below.

### Bottom-up mass spectrometry of protein samples

All protein samples were incubated with 7.5 M urea for 15 min at 37 °C, then reduced with 4 mM TCEP for 20 min at 37 °C. Chloroacetamide was added to a final concentration of 12 mM and the samples were incubated for 15 min at 37 °C. 1.5 ng/μL endoproteinase LysC was added and the digestion proceeded for 1 hour at 37 °C. Samples were diluted to 2 M urea with 50 mM HEPES, (pH 8.0) and a final concentration of 1 mM CaCl_2_ final concentration was added. Trypsin was added at a concentration of 0.5 ng/μL and incubated overnight at 37 °C. Samples were desalted using Pierce™ C18 Spin Tips & Columns (ThermoFisher Scientific, product #89870). After desalting and drying, peptides were suspended in water containing 0.2% formic acid and 2% acetonitrile for further LC-MS/MS analysis. LC-MS/MS analysis was performed with an EASY-nLC 1200 (ThermoFisher Scientific, San Jose, CA) coupled to a Q Exactive HF hybrid quadrupole-Orbitrap mass spectrometer (ThermoFisher Scientific, San Jose, CA). Peptides were separated on an Aurora UHPLC Column (25 cm × 75 μm, 1.6 μm C18, AUR2-25075C18A, Ion Opticks) with a flow rate of 0.35 μL/min for a total duration of 75 min and ionized at 1.6 kV in the positive ion mode. The gradient was composed of 6% solvent B (2 min), 6-25% B (20.5 min), 25-40% B (7.5 min), and 40–98% B (13 min); solvent A: 2% acetonitrile (ACN) and 0.2% formic acid (FA) in water; solvent B: 80% ACN and 0.2% FA. MS1 scans were acquired at the resolution of 60,000 from 375 to 1500 m/z, AGC target 3e6, and maximum injection time 15 ms. The 12 most abundant ions in MS2 scans were acquired at a resolution of 30,000, AGC target 1e5, maximum injection time 60 ms, and normalized collision energy of 28. Dynamic exclusion was set to 30 s and ions with charge +1, +7, +8 and >+8 were excluded. The temperature of ion transfer tube was 275 °C and the S-lens RF level was set to 60. MS2 fragmentation spectra were searched with Proteome Discoverer SEQUEST (version 2.5, Thermo Scientific) against *in silico* tryptic digested Uniprot all-reviewed *Azotobacter vinelandii* database. The maximum missed cleavages were set to 2. Dynamic modifications were set to oxidation on methionine (M, +15.995 Da), deamidation on asparagine and glutamine (N and Q, +0.984 Da) and protein N-terminal acetylation (+42.011 Da). The maximum parental mass error was set to 10 ppm, and the MS2 mass tolerance was set to 0.03 Da. The false discovery threshold was set strictly to 0.01 using the Percolator Node validated by q-value. The relative abundance of parental peptides was calculated by integration of the area under the curve of the MS1 peaks using the Minora LFQ node.

### Cryo-EM sample preparation and data collection

For all samples, three to four μL of sample were applied to freshly glow-discharged Quantifoil R1.2/1.3 300 mesh ultrathin carbon grids and blotted for 1-3 seconds with a blot force of 6, at ∼90% humidity using a Vitrobot Mark IV (FEI) in an anaerobic chamber ^44^. In detergent containing conditions (Pgi1, GlnA), equivalent volumes of a 1% (*w*/*v*) CHAPSO stock were mixed with the protein solution immediately before application to the grid. Grids were stored in liquid nitrogen until data collection. Datasets were collected with a 6k x 4k Gatan K3 direct electron detector and Gatan energy filter on a 300 keV Titan Krios in superresolution mode using SerialEM ^45^ at a pixel spacing of 0.325 Å. A total dose of 60 e^-^/Å^2^ was utilized with a defocus range of -0.8 to -3.0 μm at the Caltech Cryo-EM Facility.

### Cryo-EM image processing

Processing of datasets was performed in cryoSPARC 4.4.1^4^. Movie frames were aligned and summed using patch motion correction, and the contrast transfer function (CTF) was estimated using the patch CTF estimation job in cryoSPARC. Templates for template picking or filament tracing in cryoSPARC were generated by manually picking from a subset of micrographs, in the case of Bfr blob picking with a circular template was used. Picked particles were used for multiple rounds of reference-free 2D classification, *ab initio* model generation, and heterogeneous refinement (**Supp. Fig. 1**). Global CTF refinement was carried out, followed by non-uniform refinement^4^. Resolutions were estimated by the gold-standard Fourier shell correlation (FSC) curve with a cut-off value of 0.143. Three dimensional variability analysis^5^ of Pgi1 was carried out on 2x downsampled particles following Non-Uniform Refinement in cryoSPARC 4.4.1 with 3 modes over 20 iterations using a filter resolution of 5 Å. Maps representing variability components were calculated using ‘intermediate’ mode with 5 frames and no filters. These maps were re-extracted with no binning and put through homogeneous refinement.

### CryoET sample preparation

*A. vinelandii* cells were grown to ∼1.2 OD_600_ in nitrogen-free Burk’s media. 3 mL of culture were pelleted at 5,000g for 3 min. The cell pellet was resuspended in 130 μL of fresh nitrogen-free Burk’s media. EM grids (Quantifoil R2/2 Cu 200 mesh) were glow dischared using an Emitech K100X. Samples were prepared via manual blotting between each of 4 applications of 3 μL of the concentrated cell culture within a Vitrobot Mark IV (ThermoFisher Scientific) equilibrated to 95% relative humidity at 6°C.

Cryogenic focused ion beam (cryoFIB) milling was performed using an Aquilos II cryo-FIB/SEM microscope (ThermoFisher Scientific). Targets with appropriate ice thickness for milling were selected on the grid. A platinum layer (∼10 nm) was sputter coated and a gas injection system (GIS) was used to deposit the precursor compound trimethyl(methylcyclopentadienyl) platinum (IV). The stage was tilted to 12°, corresponding to a milling angle of 15° relative to the plane of grids. FIB milling was performed using stepwise decreasing current as the lamellae became thinner (3.0 nA to 50 pA, final thickness: ∼160 nm). The grids were then stored in liquid nitrogen before data collection.

### CryoET data collection and processing

Tilt series were collected on a 300-kV Titan Krios transmission electron microscope (Thermo Fisher Scientific) equipped with a high brightness field emission gun (xFEG), a spherical aberration corrector, a Bioquantum energy filter (Gatan), and a K3 Summit detector (Gatan). The images were recorded at a nominal magnification of 42,000x in super-resolution counting mode using SerialEM. For each tilt series, images were acquired using a dose-symmetric scheme between -48° and 48° relative to the lamella with 3° increments. At each tilt angle, the image was recorded as movies divided into eight subframes. The total electron dose applied to a tilt series was 150 e-/Å^2^. The defocus target was set to be -2 μm. All movie frames were corrected with a gain reference collected in the same EM session.

Movement between frames was corrected using MotionCor2 without dose weighting. All tilt series were aligned using AreTomo and the tomograms were inspected to identify good tilt series. Segmentation was performed using Dragonfly software, Version 2024.1 Build 12611 for Linux.

### Model building and refinement

Initial models were generated from AlphaFold2 predictions^12^. Multiple iterations were carried out of manual model building and ligand fitting in Coot^46^, and refinement in Refmac5^47^ and Phenix^48^. Data collection, refinement, and validation are presented in Supplementary Table 5.

## Data and materials availability

The single particle cryo-EM maps and models have been deposited into the PDB and EMDB for release upon publication.

## Figure preparation and presentation

Structural figures were prepared in ChimeraX and Pymol^49^. Figures were compiled in Adobe Illustrator.

## Author contributions

R.A.W. designed experiments. R.A.W., A.O.M., and Y.S. performed sample preparation, biochemical analyses, electron microscopy data collection and analyses, and model building and refinement. T.Z. assembled curated AlphaFold databases. R.A.W. supervised all research. R.A.W. wrote the manuscript, and all authors contributed to revisions.

## Acknowledgments

This work was funded by support from the Howard Hughes Medical Institute, NIH 1F32GM143836 (R.A.W.), and NIH 1K99GM152765 (R.A.W.). We thank Dr. Douglas Rees, Welison Floriano, Dr. Songye Chen, and Dr. Julian Braxton for invaluable discussions. Bottom-up mass spectrometry of protein samples was performed at the Beckman Proteome Exploration Laboratory supported by the Arnold and Mabel Beckman Foundation. Generous support of the Beckman Institute for the Caltech Cryo-EM Resource Center was essential for the performance of this research. We thank Momoko Shiozaki, Xiaowei Zhao, Nikki Jean, and Rui Yan at the HHMI Janelia CryoEM Facility for help in sample preparation, microscope operation, and data collection for cryoET.

