## Supplementary Information for "CryoEM-enabled visual proteomics reveals *de novo* structures of oligomeric protein complexes from *Azotobacter vinelandii*"

**Supplementary Table 1:** *Azotobacter vinelandii* strains used in this study

| **Strain** | **Relevant characterisitcs** | **Source or reference** |
|---|---|---|
| DJ605 | Δ*nifV* | Reference 28: Jacobson *et al*. |
| $MoFe^{Anc1}$ | Anc1-MoFe-Protein | This study, based on reference 19: Garcia *et al.* |
| NifH-UnaG | NifH C-terminally tagged with UnaG | This study |

| **Supplementary Table 2: Phosphoglucoisomerase**<br>Mass spectrometry results are listed in order of Score Sequest HT. | | |
|---|---|---|
| DALI vs. PDB | DALI vs. AlphaFold | Mass Spectrometry |
| Glucose-6-phosphate isomerase (PDB 4QFH) | Glucose-6-phosphate isomerase (C1DDK7) | Nitrogenase protein alpha chain OS=Azotobacter vinelandii (strain DJ / ATCC BAA-1303) OX=322710 GN=nifD PE=3 SV=1 |
| Glucose-6-phosphate isomerase (PDB 3IFS) | Glucose-6-phosphate isomerase (C1DFK1) | Nitrogenase molybdenum-iron protein beta chain OS=Azotobacter vinelandii (strain DJ / ATCC BAA-1303) OX=322710 GN=nifK PE=3 SV=1 |
| Putative glucosamine-fructose-6-phosphate aminotr (PDB 2A3N) | Glucose-6-phosphate isomerase (C1DSC1) | Glucose-6-phosphate isomerase OS=Azotobacter vinelandii (strain DJ / ATCC BAA-1303) OX=322710 GN=pgi-1 PE=3 SV=1 |
| Transcriptional regulator (PDB 4IVN) | Arabinose 5-phosphate isomerase (C1DQ58) | Phosphoenolpyruvate synthase OS=Azotobacter vinelandii (strain DJ / ATCC BAA-1303) OX=322710 GN=ppsA PE=3 SV=1 |
| Phosphoheptose isomerase (PDB 2X3Y) | Transcriptional regulatory protein, rpir family (C1DRP7) | Nitrogenase iron-molybdenum cofactor biosynthesis protein NifE OS=Azotobacter vinelandii (strain DJ / ATCC BAA-1303) OX=322710 GN=nifE PE=1 SV=1 |

| **Supplementary Table 3: Glutamine Synthetase**<br>Mass spectrometry results are listed in order of Score Sequest HT. | | |
|---|---|---|
| DALI vs. PDB | DALI vs. AlphaFold | Mass Spectrometry |
| L-Glutamine synthetase (PDB 5DM3) | Glutamine synthetase (C1DHW3) | Nitrogenase iron protein OS=Azotobacter vinelandii (strain DJ / ATCC BAA-1303) OX=322710 GN=nifH PE=1 SV=1 |
| Glutamine synthetase (PDB 6PEW) | Glutamine synthetase catalytic domain family protein (C1DQF3) | Pyruvate dehydrogenase E1 component OS=Azotobacter vinelandii (strain DJ / ATCC BAA-1303) OX=322710 GN=aceE PE=4 SV=1 |

| | | |
|---|---|---|
| Probable glutamine synthetase (PDB 4HPP) | Glutamine synthetase (C1DK83) | Nitrogenase protein alpha chain OS=Azotobacter vinelandii (strain DJ / ATCC BAA-1303) OX=322710 GN=nifD PE=3 SV=1 |
| Glutamate-ammonia ligase domain-containing protein (PDB 2J9I) | Putative glutamine synthetase (C1DK81) | Nitrogenase molybdenum-iron protein beta chain OS=Azotobacter vinelandii (strain DJ / ATCC BAA-1303) OX=322710 GN=nifK PE=3 SV=1 |
| Bestrophin (PDB 8ECY) | Transglutaminase domain protein (C1DJA3) | Glutamine synthetase OS=Azotobacter vinelandii (strain DJ / ATCC BAA-1303) OX=322710 GN=glnA PE=3 SV=1 |

**Supplementary Table 4: Filaments sample**
Mass spectrometry results are listed in order of Score Sequest HT.

| DALI vs. PDB TssC | DALI vs. PDB SthA | DALI vs. AF TssC | DALI vs. AF SthA | Mass Spectrometry |
|---|---|---|---|---|
| Inorganic polyphosphate/atp-glucomannokinase (PDB 1WOQ) | Dihydrolipoyl dehydrogenase (PDB 3L8K) | DUF877 family protein (C1DM91) | Soluble pyridine nucleotide transhydrogenase (C1DR10) | Flagellin OS=Azotobacter vinelandii (strain DJ / ATCC BAA-1303) OX=322710 GN=fliC PE=3 SV=1 |
| Transposable element p transposase (PDB 6PE2) | RV3303C-LPDA (PDB 1XDI) | Uncharacterized protein (C1DJS3) | Dihydrolipoyl dehydrogenase (C1DM54) | Uncharacterized protein OS=Azotobacter vinelandii (strain DJ / ATCC BAA-1303) OX=322710 GN=Avin_16040 PE=4 SV=1 (putative surface layer protein) |
| CRISPR-associated endonuclease, csn1 family (PDB 8D2Q) | FAD-dependent pyridine nucleotide-disulphide (PDB 3NT6) | Phage p2 tail sheath fi-like protein (C1DS10) | Glutathione-disulfide reductase (C1DIK4) | DUF877 family protein OS=Azotobacter vinelandii (strain DJ / ATCC BAA-1303) |

| | | | | |
|---|---|---|---|---|
| | | | | OX=322710 GN=Avin_50810 PE=4 SV=1 |
| Putative kinase (PDB 3LM2) | NADPH oxidase (PDB 2CDU) | Periplasmic substrate-binding protein of sugar ab (C1DLB0) | Assimilatory nitrite reductase (C1DHD0) | Soluble pyridine nucleotide transhydrogenase OS=Azotobacter vinelandii (strain DJ / ATCC BAA-1303) OX=322710 GN=sthA PE=3 SV=1 |
| CRISPR-associated endonuclease, csn1 family,crisp (PDB 6WBR) | Thioredoxin glutathione reductase (PDB 2X8C) | Periplasmic substrate-binding protein of sugar ab (C1DLB1) | fFAD-dependent pyridine nucleotide-disulphide (C1DJB9) | 60 kDa chaperonin OS=Azotobacter vinelandii (strain DJ / ATCC BAA-1303) OX=322710 GN=groL PE=3 SV=1 |

| Datasets | Pgi1 | GlnA | TssC | SthA | FliC | Bfr |
|---|---|---|---|---|---|---|
| PDB IDs | 9N4W | 9N4X | 9N4V | 9N4Y | 9N59 | 9N5A |
| Microscope | Titan Krios | Titan Krios | Titan Krios | Titan Krios | Titan Krios | Titan Krios |
| Camera | Gatan K3 Summit | Gatan K3 Summit | Gatan K3 Summit | Gatan K3 Summit | Gatan K3 Summit | Gatan K3 Summit |
| Magnification | 130,000x | 130,000x | 130,000x | 130,000x | 130,000x | 130,000x |
| Voltage (kV) | 300 | 300 | 300 | 300 | 300 | 300 |
| Recording mode | counting | counting | counting | counting | counting | counting |
| Frames/Movies | 40 | 40 | 40 | 40 | 40 | 40 |
| Total Electron dose (e-/Å2) | 60 | 60 | 60 | 60 | 60 | 60 |
| Defocus range (µm) | -0.8 to -3.0 | -0.8 to -3.0 | -0.8 to -2.5 | -0.8 to -2.5 | -0.8 to -2.5 | -0.8 to -2.5 |
| Pixel size (Å) | 0.65 | 0.65 | 0.65 | 0.65 | 0.65 | 0.65 |
| Micrographs collected | 16,547 | 6,280 | 10,154 | 10,154 | 10,154 | 10,154 |
| Micrographs used | | | | | | |
| Total extracted particles | 10,772,123 | 3,449,407 | 617,466 | 5,140,089 | 617,466 | 6,429,184 |
| Refined particles | 70,453 | 10,672 | 265,571 | 88,454 | 47,453 | 15,105 |
| Symmetry imposed | D5 | D6 | C6 | C1 | C1 | O |
| Nominal Map Resolution (Å) | 2.49 | 3.18 | 1.85 | 3.70 | 2.82 | 2.96 |
| FSC threshold | 0.143 | 0.143 | 0.143 | 0.143 | 0.143 | 0.143 |
| masked/unmasked | 2.4/2.5 | 3.1/3.2 | 1.8/1.8 | 3.7/3.7 | 2.8/3.1 | 2.9/2.9 |
| Refinement | | | | | | |
| Number of atoms | | | | | | |
| Protein | 43,250 | 39,859 | 88,086 | 50,696 | 8,088 | 30,804 |
| Ligand | | | | FAD:14 | | HEM:12 |
| MapCC (mask/box) | 0.86/0.74 | 0.85/0.74 | 0.91/0.67 | 0.89/0.81 | 0.84/0.27 | 0.82/0.57 |
| Map sharpening B-factor | 83 | 54 | 52 | 65 | 45 | 99 |
| R.m.s. deviations | | | | | | |
| Bond lengths (Å) | 0.004 | 0.003 | 0.004 | 0.002 | 0.003 | 0.003 |
| Bond angles (º) | 0.545 | 0.569 | 0.614 | 0.485 | 0.561 | 0.676 |
| MolProbity score | 1.62 | 1.87 | 1.00 | 1.43 | 1.81 | 1.56 |
| Clashscore (all atom) | 8.07 | 12.54 | 2.20 | 7.79 | 9.13 | 7.65 |
| Rotamer outliers (%) | 1.26 | 0.16 | 0.45 | 0.11 | 0.00 | 1.55 |
| Ramachandran plot | | | | | | |
| Favored (%) | 97.46 | 96.16 | 98.33 | 98.00 | 95.37 | 99.35 |
| Allowed (%) | 2.54 | 3.84 | 1.67 | 1.86 | 4.63 | 0.65 |
| Outliers (%) | 0.05 | 0.00 | 0.00 | 0.14 | 0.00 | 0.00 |

**Supplementary Table 5: Cryo-EM data collection, refinement, and validation statistics**

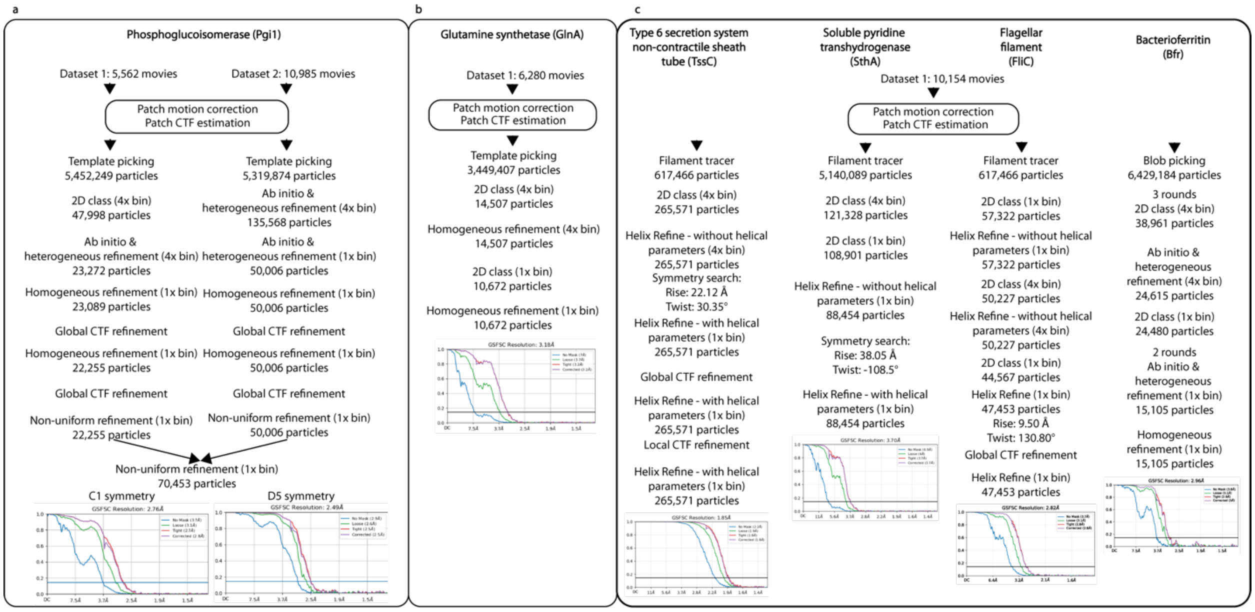


**Supplementary Figure 1: CryoEM data processing pipelines for all contaminants.** A, Processing for phosphoglucoisomerase (Pgi1). B, Processing for glutamine synthetase (GlnA). C, Processing for the filaments dataset, from left to right: Type 6 secretion system non-contractile sheath tube (TssC); the soluble pyridine transhydrogenase (SthA); the flagellar filament (FliC); and bacterioferritin (Bfr). Processing for all datasets was completed in cryoSPARC v4.4.1. Workflows are presented from top to bottom.

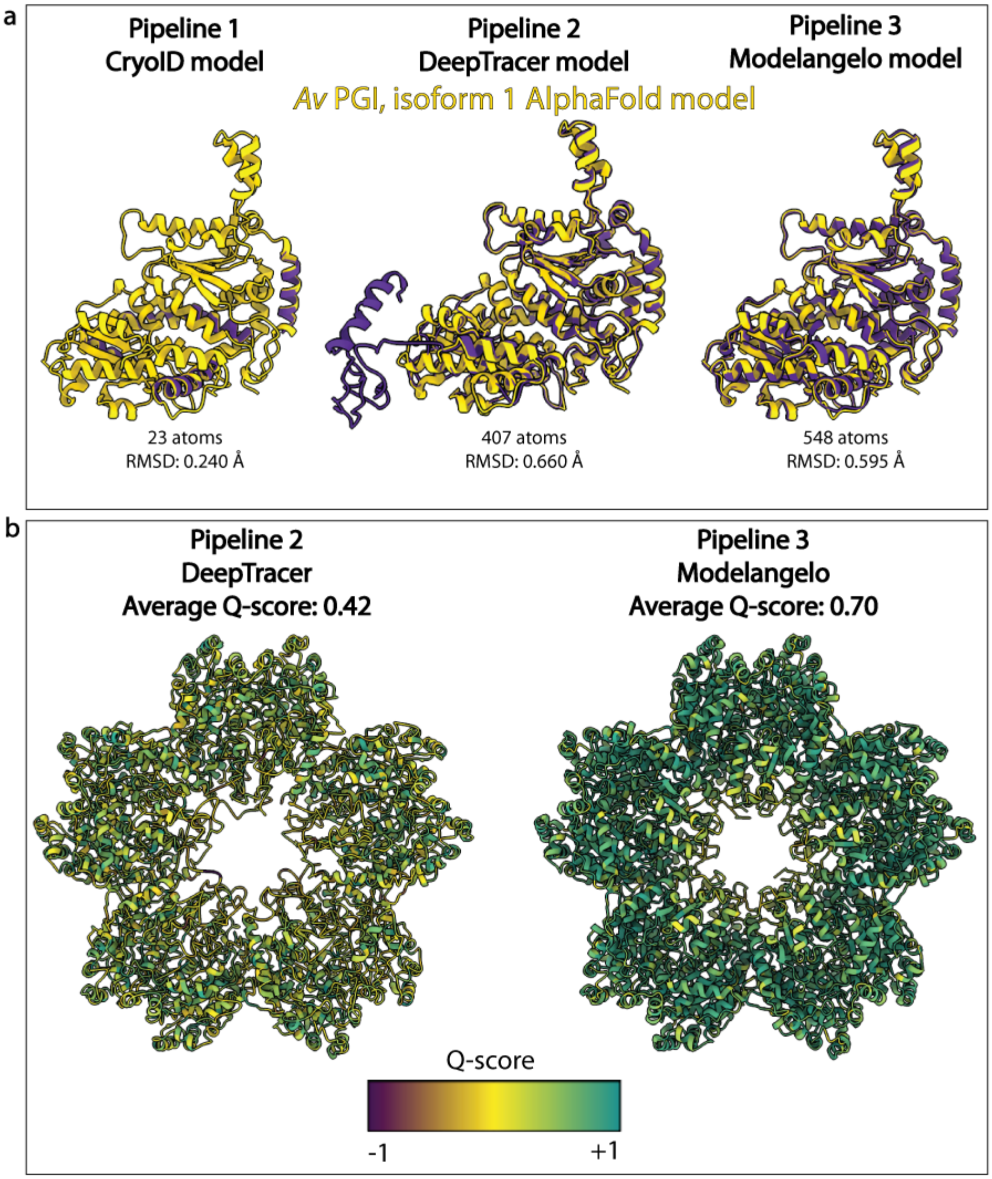


**Supplementary Figure 2: Comparison of results from three computational pipelines for the identification of PGI.** A, Alignment of models built by CryoID, DeepTracer, and ModelAngelo with the AlphaFold2 predicted model for *A. vinelandii* PGI, isoform 1 downloaded from the AlphaFold database. B, Models built by DeepTracer, and ModelAngelo color-coded according to their map-to-model fit (Q-score).

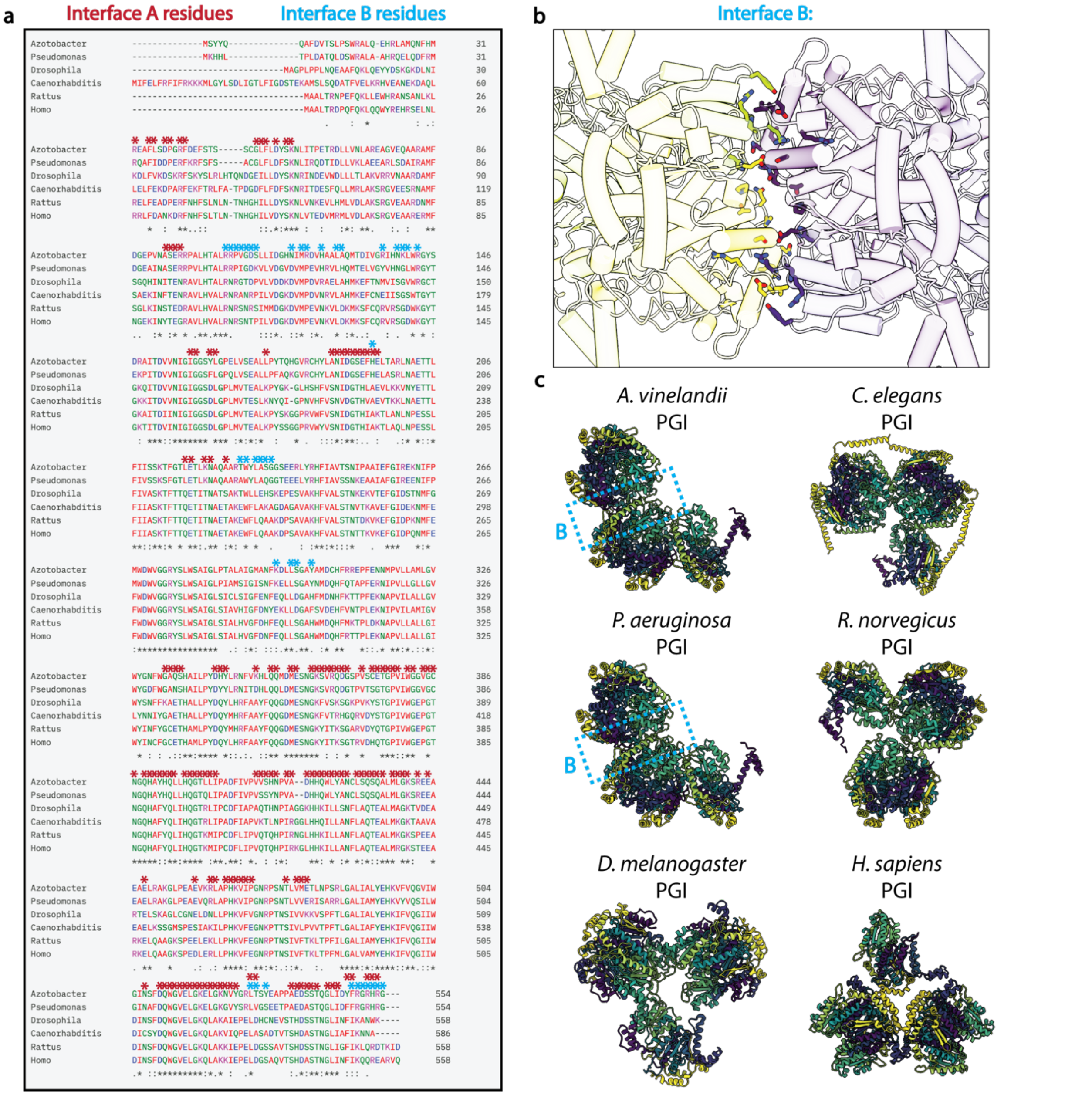


**Supplementary Figure 3: Conservation of Pgi1 interfaces across model organisms.** A, CLUSTAL sequence alignment of Pgi1 across *A. vinelandii, P. aeruginosa, Drosophila melanogaster, Caenorhabitis elegans, Rattus norvegicus,* and *Homo sapiens.* Residues are color coded according to the CLUSTAL color scheme. Residues within Interface A are marked with a red asterisk, while residues in Interface B are marked with a blue asterisk. B, Residues involved in Interface B are highlighted within the *A. vinelandii* cryoEM structure. C, Comparison of the AlphaFold3 structural predictions of the sequences shown in panel A, using 5 copies of each chain.

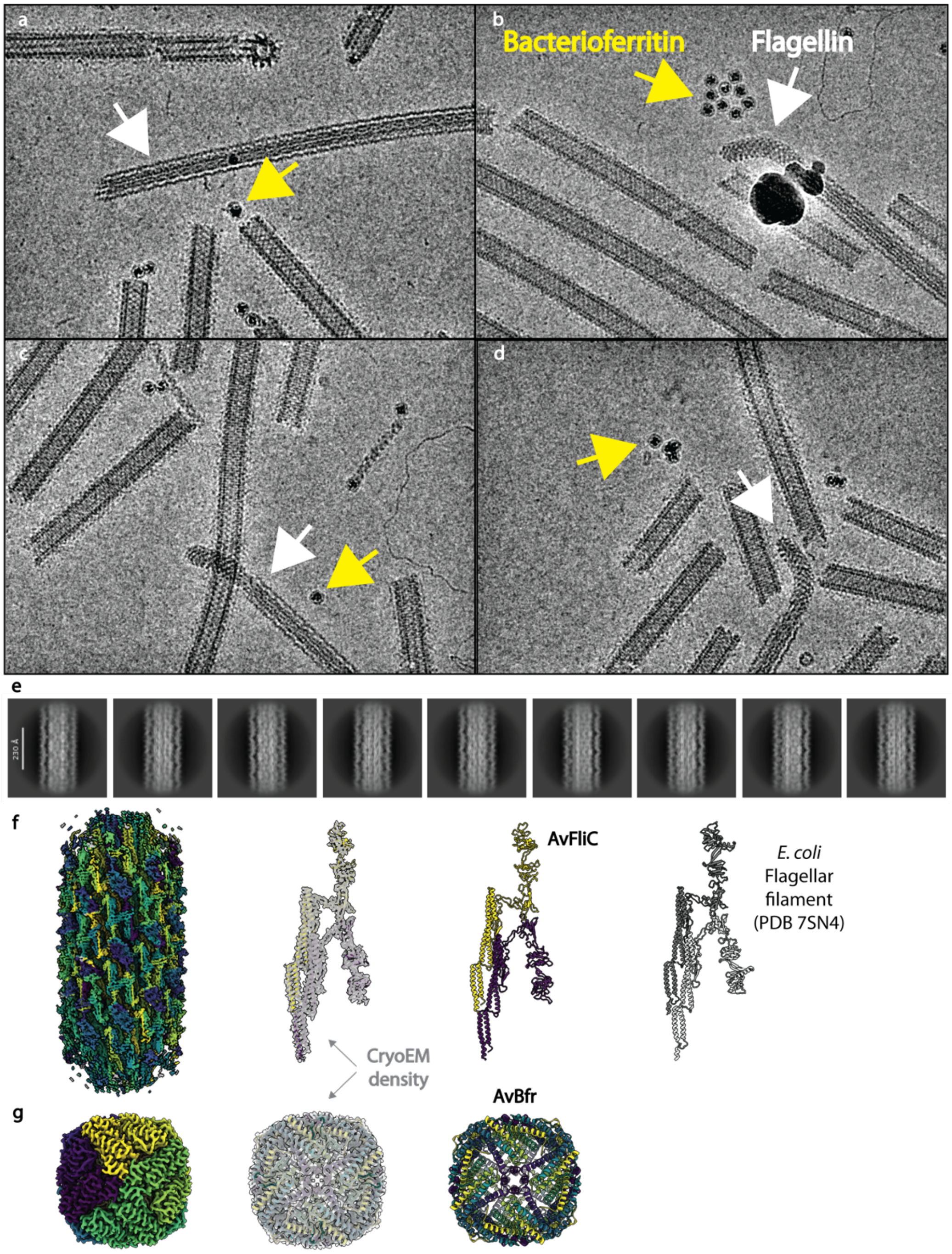


**Supplementary Figure 4: Filament C is the *Azotobacter vinelandii* flagellin filament.** A-D, Representative micrographs from the same dataset containing Filaments A and B showing "Filament C", confirmed by visual analysis and mass spectrometry to be the flagellin filament. The flagellar hook is clearly seen in panels B-D. Yellow arrows indicate bacterioferritin molecules. E, 2D class averages of the flagellin filament. F, 2.82 Å resolution reconstruction of the flagellar filament compared with the structure of the *E. coli* flagellar filament. G, 2.96 Å resolution reconstruction of bacterioferritin.

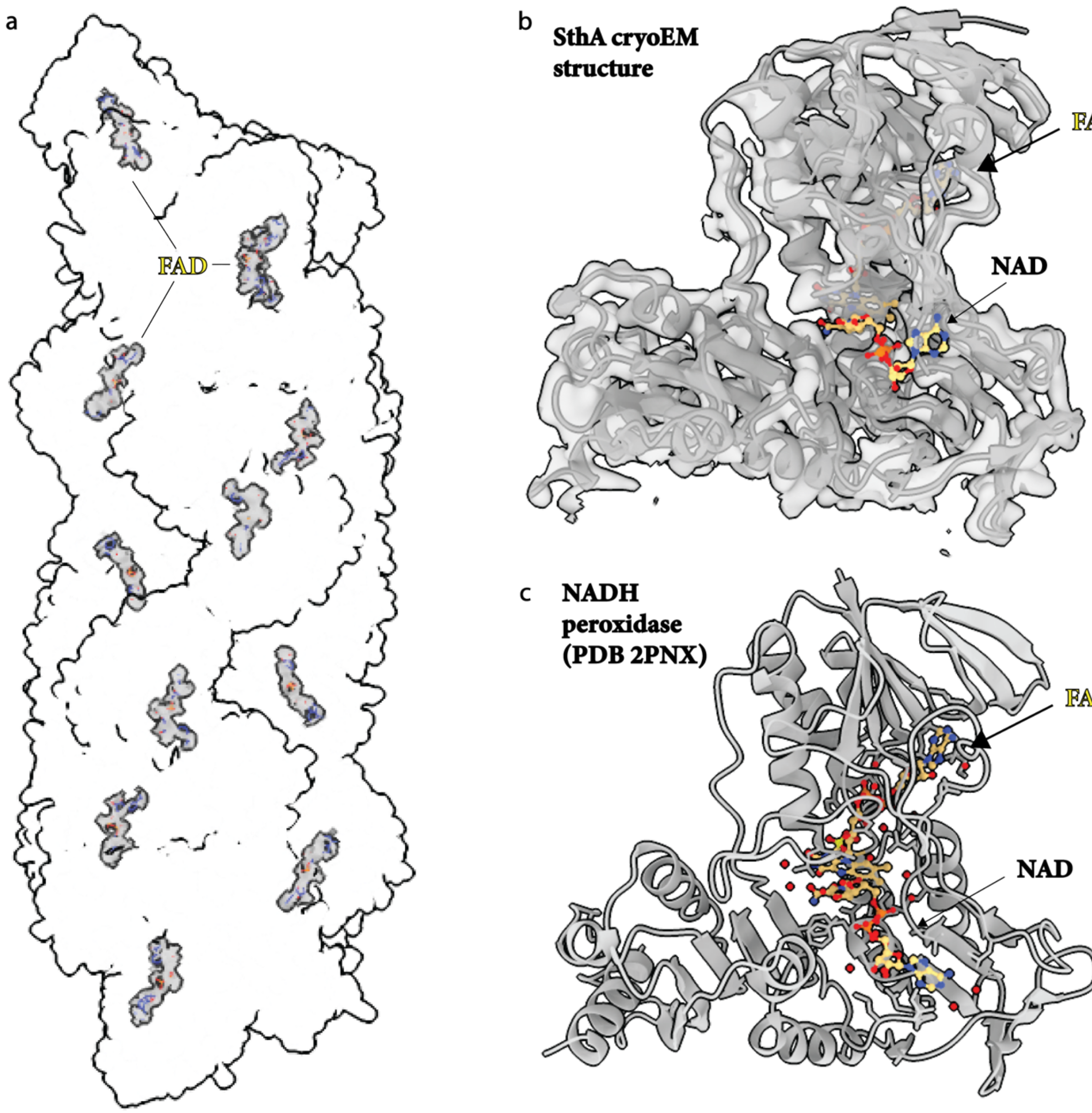


**Supplementary Figure 5: Cofactor and substrate binding within the soluble pyridine transhydrogenase (SthA) filament.** A, CryoEM density surrounding the FAD cofactor is shown in gray throughout the SthA filament. B, CryoEM density and model for one SthA monomer with the AlphaFold-predicted model for bound NAD shown in yellow. No cryoEM density (gray) was observed at this site. C, Structure of a homologous NADH peroxidase with FAD and NAD bound.
